# An RNA origami robot that traps and releases a fluorescent aptamer

**DOI:** 10.1101/2023.05.19.541473

**Authors:** Néstor Sampedro Vallina, Ewan K.S. McRae, Cody Geary, Ebbe Sloth Andersen

## Abstract

RNA nanotechnology aims at using RNA as a programmable material to create self-assembling nanodevices for application in medicine and synthetic biology. RNA devices have been developed by adopting mechanisms such as allosteric binding and toehold-mediated strand displacement. There are, however, no examples of RNA “robotic” devices that sense, compute, and actuate through mechanical reconfiguration as has been demonstrated in DNA nanotechnology. Here we use the RNA origami method to prototype an RNA robotic device, named the “Traptamer”, that senses two RNA key strands, acts as a Boolean AND gate, and activates the fluorescent aptamer iSpinach through release from a mechanical trap. The Traptamer depends on binding of two different RNA key strands to achieve full activation and can be reversed by addition of two complementary RNA anti-key strands. Cryo-EM of the closed Traptamer structure at 5.45 Å resolution reveals a hinge-like mechanical distortion of the iSpinach motif. Our RNA robot prototype opens the door to build more sophisticated RNA machines that use sensing, computing, and acting modules to precisely control RNA functionalities.

## INTRODUCTION

DNA nanotechnology exploits the regular structure of DNA double helices to construct higher-order 3D structures and the simple base pairing rules to design sequences that self-assemble into these structures [1]. To create dynamic structures, the toehold-mediated strand-displacement (TMSD) mechanism is often used to reconfigure base pairings to create molecular machines [2, 3] or logic circuits [4-6]. The DNA origami method has paved the way for the creation of intricate DNA nanostructures through the heat-annealing of a long scaffold strand with numerous smaller staple strands [7]. With the advent of DNA origami, it became possible to engineer more complex molecular machines, including a DNA origami box that incorporated two TMSD locks that responded as an AND gate to two specific key signals to open the device [8]. This nanomechanical device can be referred to as “nanorobot”, since it can sense inputs, do basic computations, and operate within their nanoscale environment. Several examples of these robotic devices have been developed to solve tasks of targeting and killing cancer cells [9], control the accessibility of an enzyme to the substrate [10], deliver immunostimulatory cargoes to dendritic cells [11] and blood clotting agents to tumors [12].

RNA nanotechnology developed independently by using the wider variety of structural and functional motifs found in natural RNAs [13]. Tertiary RNA motifs consist of both canonical and non-canonical base pairs, which renders the base pairing rules more intricate and the design of suitable sequences more challenging [14, 15]. Several RNA nanoparticles have been developed by the tecto method and used for a variety of applications [16-20]. The RNA origami method was developed by combining crossover motifs and tertiary loop motifs to form RNA nanostructures that fold from a single strand [21]. The structures were shown to be able to self-assemble by the process of cotranscriptional folding, an important feature that enables the structures to be genetically encoded and expressed in cells [22]. The recently developed RNA origami design software allows large structures to be designed with addition of functional motifs [23]. However, the RNA origami method has mainly been used to create functional scaffolds for RNA medicine [24, 25] and RNA synthetic biology [26, 27] and structural characterization by cryo-EM [28-30]. RNA origami has been used to create dynamic devices that uses FRET between fluorescent aptamers to sense small conformational changes of target binding aptamers or larger conformational changes upon TMSD in branched kissing loops [22], but has not yet been used for implementing robotic devices that both sense, compute, act.

Inspiration for the rational design of RNA devices can be gathered from the mechanisms of riboswitches which are widely found in nature [31-33]. Riboswitches work by base pair reconfiguration induced by ligand binding, which triggers the folding and activation of a specific functional motif [34, 35]. Inspired by these mechanisms, researchers have combined target binding aptamers with functional sequences to control activities such as splicing [36], transcription [37] and translation [38, 39]. Alternatively, researchers have used toehold switches that through strand displacement activate gene expression [39]. By combining more RNA devices to control one gene, it is possible to implement Boolean logic in the activation or deactivation of expression [38-41]. Development of fluorescent aptamer technology provided RNA devices with a fluorescence output signal [42, 43]. By connecting a ligand binding aptamer with the fluorescent aptamer through a transducer module, several RNA sensors for a variety of ligands were developed [44-47]. Alternatively, TMSD can be used for controlling fluorescent aptamers [48-51]. In the above examples of riboswitch and TMSD devices, the transduction mechanism works by base pair reconfiguration. However, RNA origami devices that work by nanomechanical transduction and computation to control functional RNA motifs has not yet been explored.

Here we use the RNA origami method to develop an RNA robotic device, named the “Traptamer”. This device can sense two specific RNA “key” strands, function as a Boolean AND gate, and trigger the activation of the iSpinach fluorescent aptamer by releasing it from a mechanical trap. Several designs were screened to obtain the desired trapping of the iSpinach aptamer and fluorescence suppression. Branched kissing loops (bKL) [52] with stem loops are used as points of invasion of RNA key strands. Two RNA strands act as “keys” to open the bKL interactions and thus release the iSpinach aptamer to fold into its active conformation, bind DFHBI-1T, and fluoresce. The Traptamer is shown to activate in 20 min depending on the concentration of K^+^ and key strands. By extending the invader strands with toeholds, we demonstrate the reversibility of the trapping mechanism by removing the invaders with anti-invaders. Cryo-EM of the closed Traptamer structure at 5.45-Å resolution reveals the mechanical mode of distortion of the iSpinach motif. Thus, we have demonstrated an RNA robotic device that uses a novel mechanical distortion mechanism to control RNA functionality. The generality of the approach may be applicable to many RNA functional motifs and allow a wide range of regulatory control to be implemented in RNA medicines and RNA regulatory elements.

## RESULTS

### Design of an RNA origami trap for aptamer folding

RNA origami structures are composed of multiple helices that are connected by double crossovers, where the distance between crossovers are an integer number of full turns of the A-form helix of 11 base pairs (bp) per turn. However, by making some helices longer and others shorter we can introduce out-of-plane bending in the longer helices. If placing functional RNA motifs in the long helix, the motif might thus be bent out of shape to inactivate its function. We here use this concept to trap an aptamer in an RNA origami frame and refer to this device as a “Traptamer”. The RNA origami device is trapped during cotranscriptional folding resulting in a misfolded fluorescent aptamer that is incapable of activating its fluorogenic dye.

To demonstrate this concept, we placed the iSpinach aptamer in the middle helix of a 3-helix RNA origami tile, with the outer helices being short to force the iSpinach aptamer out of plane and into a deactivated conformation (Fig. 1A,B). To open the trap, we installed branched kissing loops (bKLs) [52] in the top and bottom helices. The bKLs were extended with a 5-bp stem capped by a 6-nucleotide (nt) loop, which can serve as a toehold for strand displacement, as previously demonstrated in Jepsen *et al*. [22]. When the cognate RNA strands, which we refer to as “RNA keys”, invade the bKL, the kissing-loop interaction is broken, which removes the structural constraint on the central helix and allows iSpinach back to its native state and bind the fluorogenic dye (Fig. 1C).

**Figure 1.**
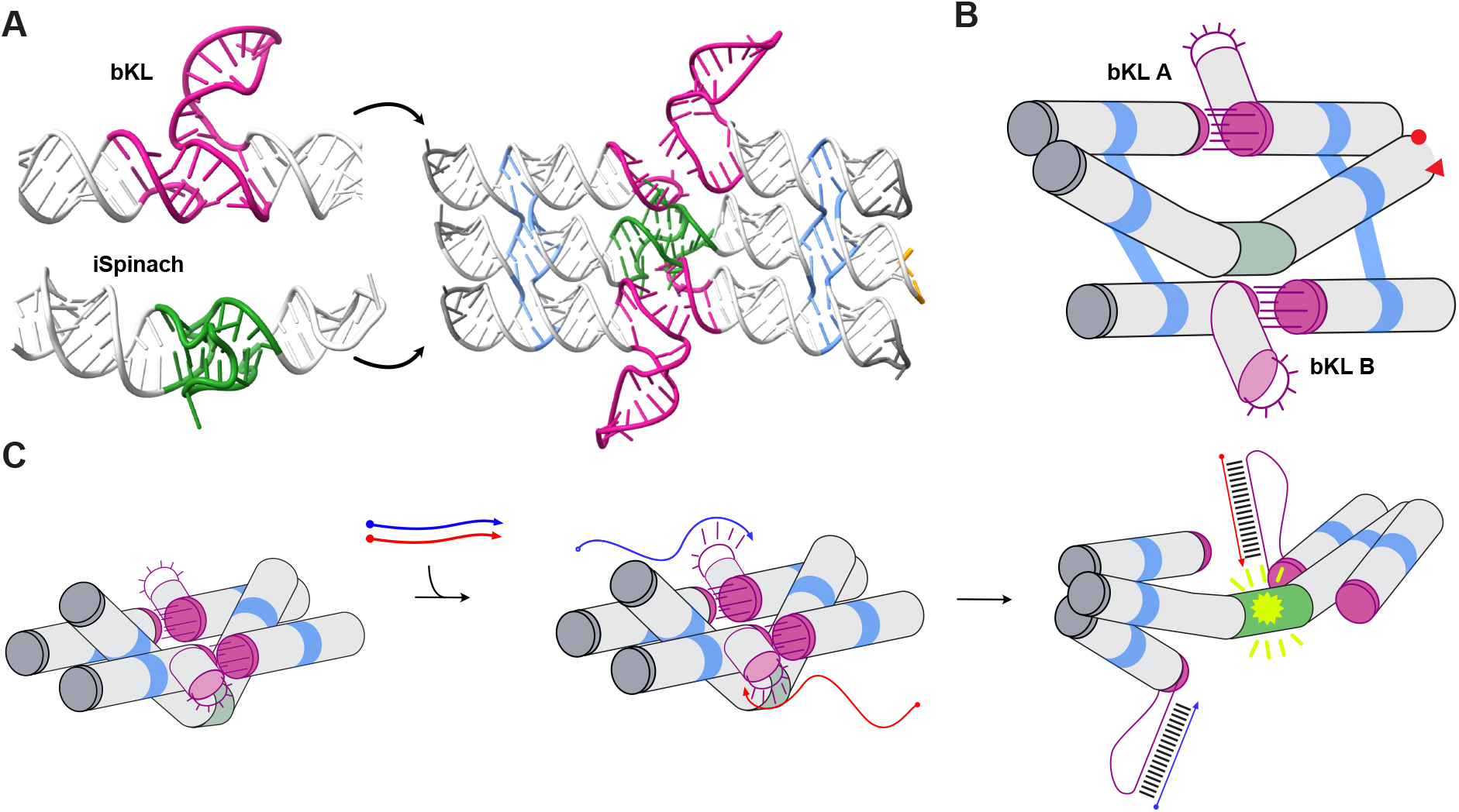
Design of a mechanical trap for aptamers. **A**. Depiction of the trap design. A 3-helix tile RNA origami bridged together by two double crossovers (blue) has the iSpinach fluorogenic aptamer (PDB: 5OB3) incorporated into the middle helix (green). The structure is bridged together by two branched kissing loops (bKLs) [52] (magenta), which have an extended 5 bp stem with a 6 nt lop at the end acting as a toehold for strand displacement. **B**. Schematic representation of the device. **C**. Schematic representation of the opening mechanism. The strand displacement reaction starts on the 5’ end of the toehold loop of the bKLs, continues along the stem of the bKL and ends up base pairing along the loop, breaking the kissing interaction, iSpinach is then allowed to fold into its active conformation and bind DFHBI-1T, producing a fluorescent output.

### Screening for iSpinach trapping and release

To identify a design with the required functionality, we generated a series of designs by varying the stem lengths around the aptamer. Each RNA design was designated “L-R”, where L and R represent the number of bp in the stems to the left and right of the iSpinach motif, respectively (Fig. 2A, Table S1). The different RNAs were produced by *in vitro* transcription, purified by size-exclusion chromatography and their fluorescence in complex with DFHBI-1T measured (see Materials and Methods). Clear differences in fluorescence were observed for the series as compared to a control where the bKL interactions were broken by tetraloops (No-KL normalized to 1.0). Designs 11-11, 12-12 and 13-12 show high fluorescence (>50%), 14-15 and 15-16 show medium fluorescence (20-50%) and 13-13, 13-14, 14-14, 15-15, 16-16 and 16-17 show low fluorescence (<20%) (Fig. 2B and Fig. S1).

**Figure 2.**
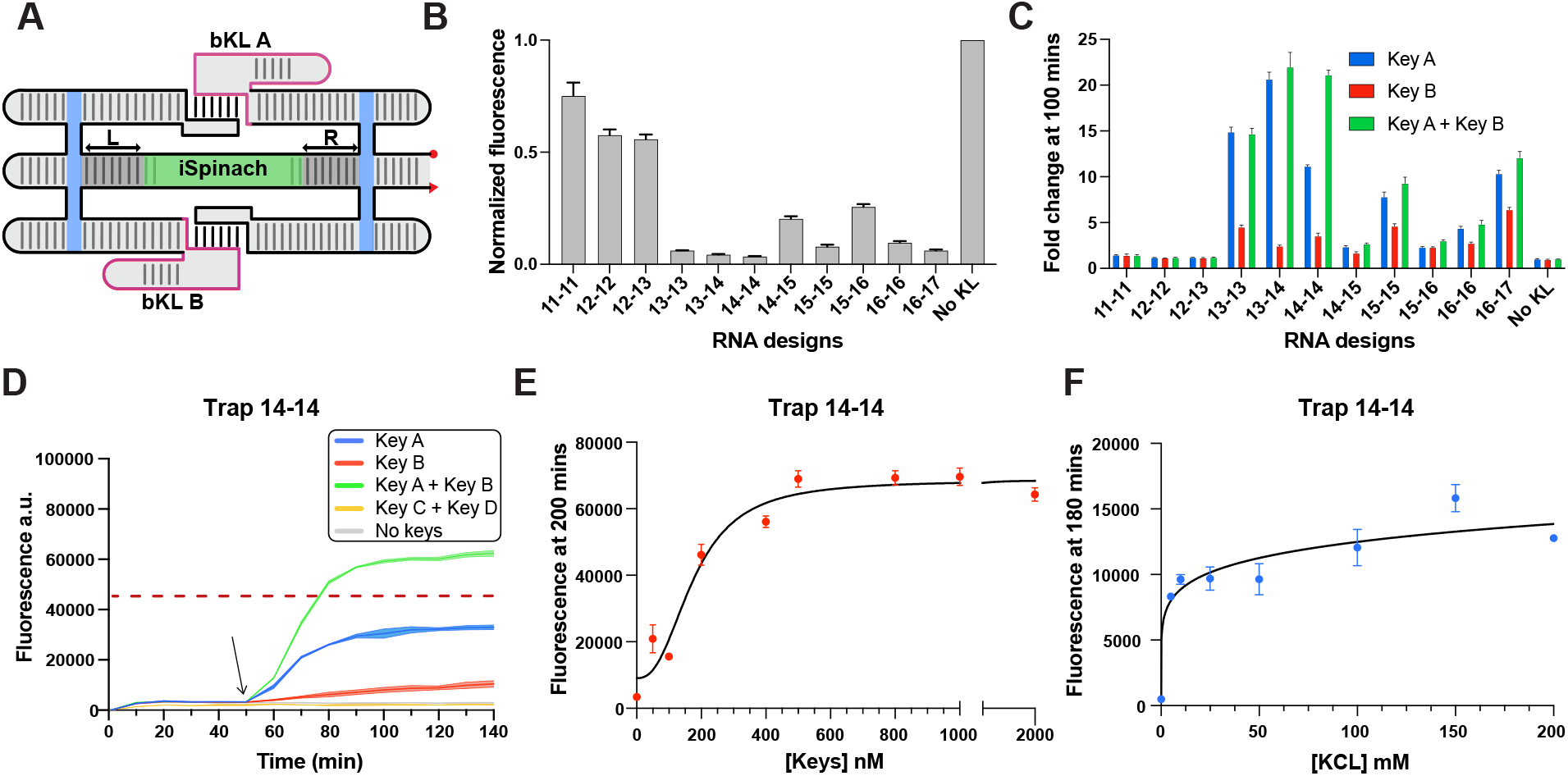
Design screening of RNA origami trapping and release. **A**.Schematic of the design parameter variations. The length of the double stranded left (L) and right (R) stems on the middle helix was varied from 11 bp to 16 on the L side and from 11 to 17 on the R side. **B**. Background fluorescence observed for the different designs when no keys were added for 100 nM RNA and 500 nM DFHBI-1T. Data corresponds to 3 technical replicates, error bars represent mean ± SD. **C**. Fold-change fluorescence activation 50 mins after addition of the RNA keys (400 nM) to the different RNA (100 nM). Fold change was calculated by comparison to the no keys sample. Data corresponds to 3 technical replicates; error bars represent mean ± SD. **D**. Observed fluorescence for the 14-14 device over time. DFHBI-1T (500 nM) was added after the first measurement, cognate and negative control keys (400 nM) were added after 45 minutes (black arrow). Data corresponds to 3 technical replicates; error bars represent mean ± SD. **E**. ssRNA key titration. Fluorescence measured after 200 minutes upon addition of different concentrations of ssRNA keys to Trap-14-14 RNA (100 nM) in complex with DFHBI-1T (1 µM). Data corresponds to 3 technical replicates; error bars represent mean ± SD. **E**. KCL titration. Fluorescence after 180 minutes upon addition of keys at different KCl concentrations. Data corresponds to 3 technical replicates; error bars represent mean ± SD.

We then tested the response of these designs to the addition of Key-A, Key-B or both. As expected, the designs that already showed high fluorescent signal in the absence of keys (11-11, 12-12, 12-13) did not respond to the addition of keys, which indicates that iSpinach is untrapped (Fig. 2C, Fig. S1). Designs with medium fluorescent signal in the absence of keys (14-15 and 15-16) responded less than 3-fold (Fig. 2C). Designs 13-13, 13-14, 15-15, 16-16 and 16-17 are fully activated by Key-A (10-20-fold) and partly by Key-B alone (2-5-fold), indicating that the formation of bKL-A is primarily responsible for disrupting the aptamer structure in these constructs. Interestingly, 14-14 shows partial activation in the presence of either key (4-11-fold), but only shows full fluorescence activation when both RNA inputs are present (22-fold). The combined signal is larger than the sum of the individual signals indicating a cooperative effect of bKL-A and bKL-B inhibiting the active form of iSpinach (Fig. 2C, Fig. S1). A fluorescent activation of 83% was observed 30 minutes after the keys were added (Fig. 2D) and alternative key sequences did not activate fluorescence, showing specificity (Fig. 2D, Key-C and Key-D). Testing the stability over extended time periods show that the signal was stable for Key-A, but less so for Key-B, which correlates with the differences in the free energy of the branch of the bKLs (Fig. S2). Given that both input strands are necessary for full activation, we consider trap 14-14 to function as an AND logic gate with a threshold set at 60% (Fig. 2D).

### The 14-14 traptamer is K^+^-dependent and modular

The properties of the 14-14 trap were investigated further. Initially the amount of key needed to obtain full opening of the trap was tested and found to reach the full response at 5X of both keys (Fig. 2E, Fig. S3C). Secondly, the impact of K^+^ concentration was tested since monovalent cations are crucial in stabilizing G-quadruplex motifs and the RNA-fluorophore complex in iSpinach [53, 54]. The fluorescence was measured before and after addition of 5X of both keys at different KCl concentrations ([KCl]) (Fig. 2F, Fig. S3D). We observed that when the [KCl] was between 5 to 100 mM, the fluorescence level gradually increased after adding the keys, which can be explained by a slow maturation of the iSpinach structure after release from the trap. On the other hand, at higher [KCl] of 150 and 200 mM, we observed a rapid increase in fluorescence upon addition of keys. This indicates that even at high [KCl], the bKLs keep the aptamer in the off state, but once released, iSpinach more rapidly adopts its mature conformation for binding to the fluorophore.

To investigate whether the observed behavior of the trap 14-14 was affected by the specific sequence design, we designed and synthesized another sequence, trap 14-14 v2, which shared the same structural elements (i. e. bKLs, aptamer sequence and 5’- and 3-ends) but has a different scaffold sequence (Table S1). We observed a similar response between trap 14-14 and 14-14 v2 (Fig. S4), suggesting that the observed behavior was not affected by the precise sequence design, but rather attributed to the structural elements of the Traptamer. To further test the modularity of our design, we incorporated the Broccoli aptamer [55] into the scaffold sequence and observed similar behavior once again (Fig. S4, Table S1). These results indicate that our design is modular, and that different aptamers or functional motifs can be regulated using similar strategies.

The excitation and emission spectra of DFHBI-1T have been shown to vary depending on the binding aptamer [55, 56], thus even small shape changes of the fluorophore-binding pocket are expected to change the fluorescence spectral characteristics. We therefore explored whether the distinct fluorescent outputs observed from the opening of 14-14 with different keys by measuring at discrete excitation and emission wavelengths (see Materials and Methods) could also be explained by different conformations of the aptamer in the Traptamer. To test this hypothesis, we measured the excitation and emission spectra of DFHBI-1T in complex with trap 14-14 in the presence of different key combinations. However, no significant difference was detected in the spectral characteristics (Fig. S5). Thus, the different fluorescence outputs observed cannot be attributed to changes in the spectral properties of DFHBI-1T.

### The trapping mechanism is reversible

The opening of the bKL by a key has previously been shown to be reversible by the addition of an anti-key strand that binds a toehold on the key strand and removes it from the complex by strand displacement [22]. By adding keys and anti-keys it may thus be possible to open and close the Traptamer to cycle the aptamer between its on and off state (Fig. 3A). We first investigated the effect of adding a 5-nt toehold to the key strand on the 5’ or 3’ end (named t-Key and Key-t, respectively). The keys without toehold activated the fluorescence at a maximum rate of 847 units/min, while t-Key activated at a maximum rate of 443 units/min and the Key-t activated at a maximum rate of 541 units/min. The t-Key showed a delayed response but reached close to maximum fluorescence after 160 min (Fig. S6). In contrast, Key-t was further delayed and seems to stabilize at 60% of maximum fluorescence (Fig. S6). The lower efficiency of Key-t can be explained by the placement of two toeholds next to each other, which has been shown to decrease the strand displacement rates in DNA systems [57]. Even though Key-t reaches a lower overall fluorescence, it does reach its own maximum faster, and we thus chose this for further experiments.

**Figure 3.**
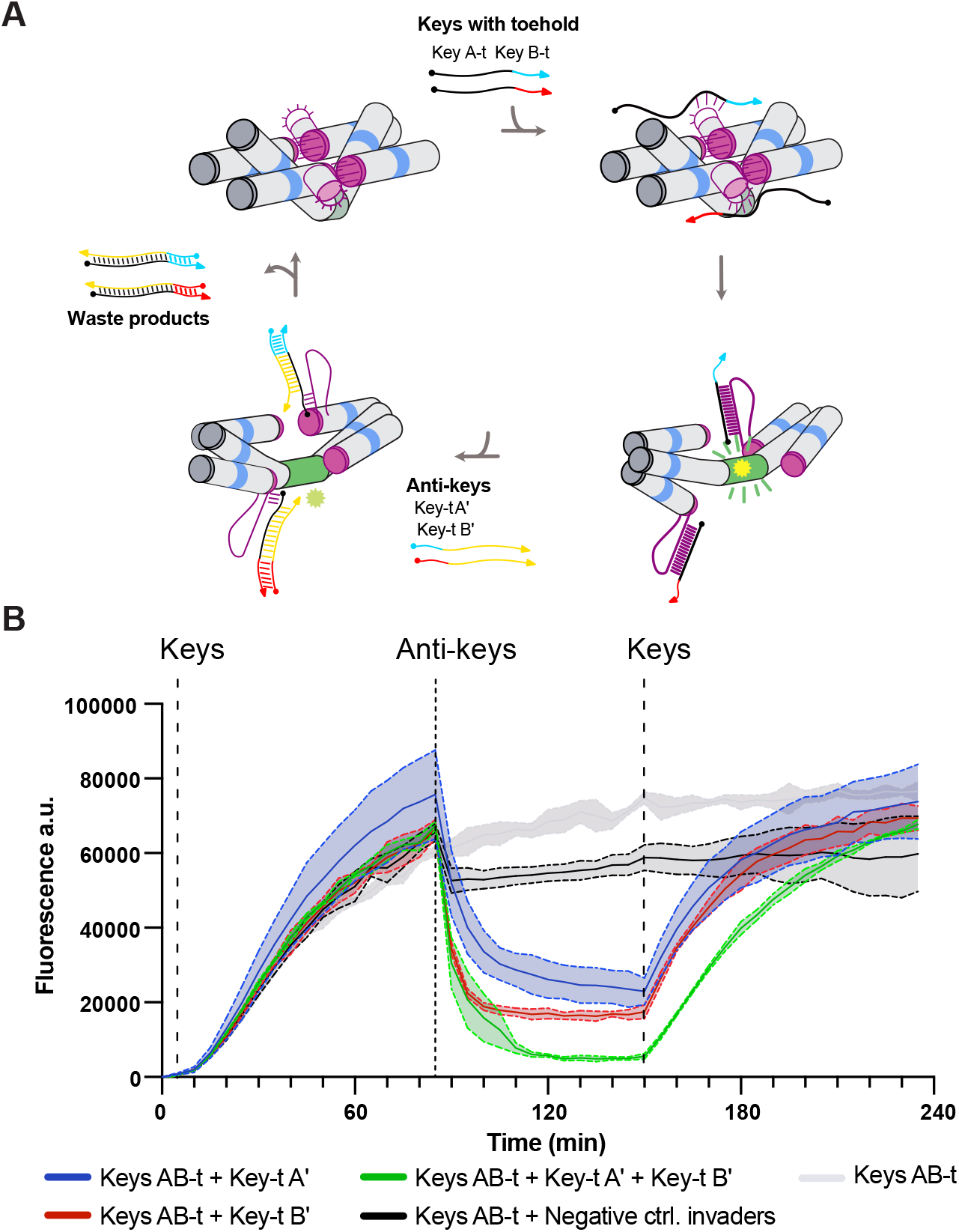
Reversible activation and deactivation of iSpinach by Trap 14-14. **A**.Schematic representation of the mechanism. The ssRNA keys were extended with a 6-nt toehold (Key-t A and Key-t B), anti-key RNA invaders were designed to displace the keys (Key-t A’ and Key-t B’). **B**. Monitored fluorescence over time of Trap 14-14 RNA (100 nM) in solution with DFHBI-1T (500 nM), keys with toehold (500 nM) were added after 5 mins (black dashed line). Key invaders (1 µM) were added after 85 mins (grey dashed line). Keys (2 µM) were added again after 150 mins Data corresponds to 3 technical replicates; error bars represent mean ± SD.

The reversibility was tested by adding 5X Key-t A and Key-t B and letting the signal increase for 80 min (Fig. 3B, left). Then a 2X excess of either one or both anti-key strands is added (Key-t A’ and Key-t B’). A fast decrease in fluorescence is observed that reaches a minimum fluorescence after 25 min (Fig. 3B, middle). It is observed that both anti-keys can turn off the fluorescent signal, with Key-t A’ being faster than Key-t B’. With both anti-keys added we observed a more efficient and closer to full deactivation. Adding keys of unrelated sequence did not lead to a significant decrease. When adding a 4X excess of Key-t A and Key-t B after 150 min, we observe a reactivation of the fluorescence that return to similar levels as the initial activation (Fig. 3B, right). The data shows that the Traptamer can trap and release the iSpinach aptamer in a reversible manner and that the activation depends on both strand displacement and maturation of the iSpinach aptamer.

### Cryo-EM reveals a hinge-like mechanical distortion of the iSpinach aptamer

The fluorescence experiments suggests that the formation of the bKLs in design 14-14 disrupts the iSpinach aptamer and prevents it from binding its fluorophore DFHBI-1T. To gain a better understanding of the trapped conformation of the iSpinach aptamer, we used cryo-EM to analyze the structure of the cotranscriptionally assembled conformation of 14-14 with and without keys. The reconstruction of the trapped conformation reached a resolution of 5.45 Å (Fig. S7, S8), and the EM map revealed well-resolved double helices and crossovers, the bKLs and a 90° bend of iSpinach (Fig. 4A,B). We also attempted to reconstruct the open state of 14-14 (i.e., after addition of the keys) with cryo-EM, but the resolution we could attain was limited by the flexibility of the molecule (Fig. S9).

**Figure 4.**
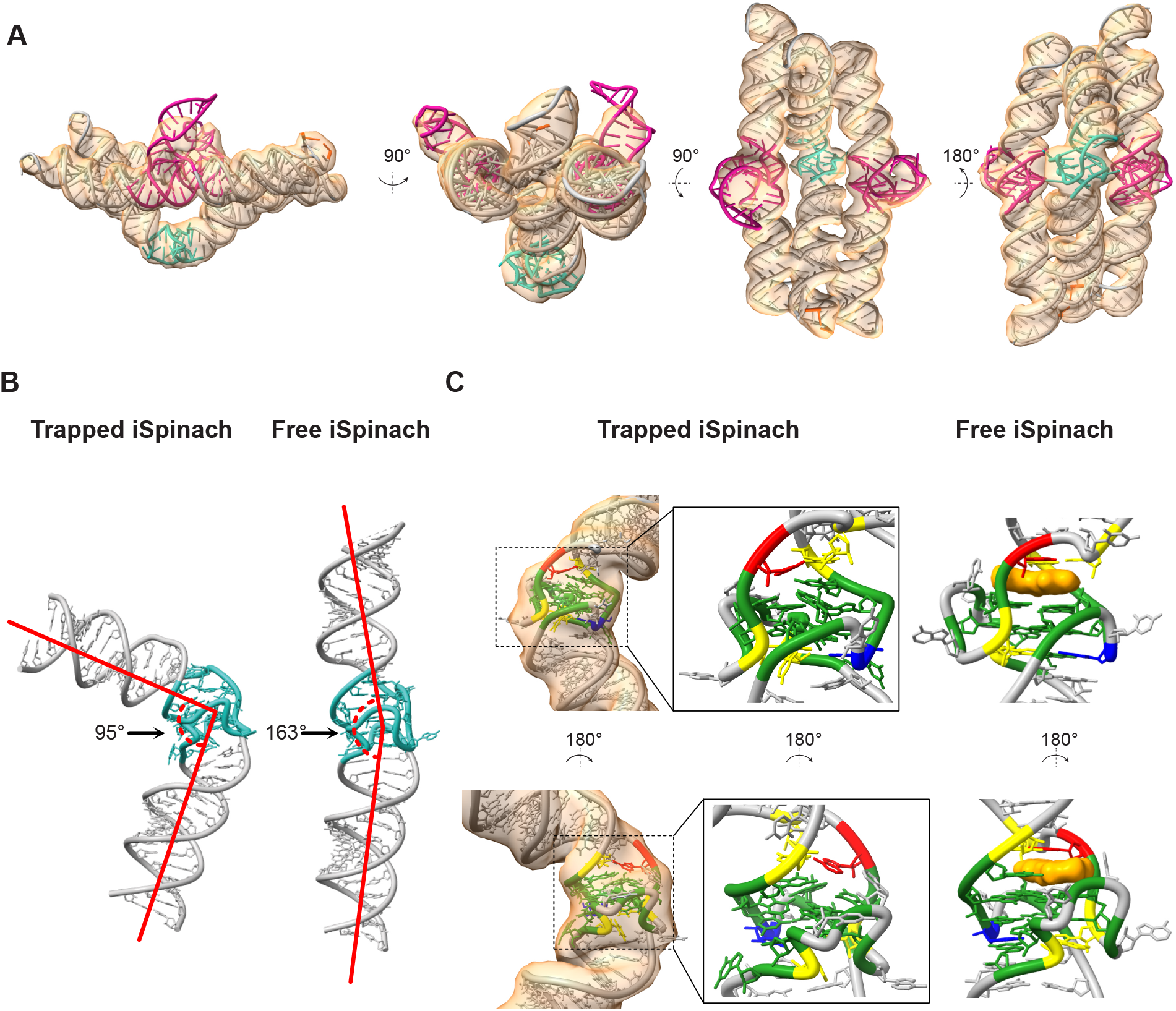
Structural characterization of the cotranscriptionally assembled state of Trap 14-14. **A**.Obtained experimental density of 5.45 Å at threshold level X (orange). The model was manually built into the density (see materials and methods), with bKLs highlighted in magenta, iSpinach in blue and the rest of the origami in grey. **B**. Comparison between the bending of the iSpinach aptamer in the traptamer (left) and a computer-generated model of iSpinach with extended helical stems (right). **C**. Comparison of the binding pocket of iSpinach. The binding pocket of iSpinach in the traptamer model appears contorted as compared to the crystal structure of iSpinach (PDB: 5OB3). Nucleotides colored as follows: G: green, C: blue, A: red, U: yellow.

To gain a deeper understanding of how iSpinach is disrupted, we manually built a model of the 14-14 trap into the density map (see Materials and Methods) (Fig. 4A and S8). By flexibly fitting the native iSpinach model into our trap 14-14 density map, we obtained a model of the deformed state and the effects on the fluorophore binding pocket (Fig. 4C). In the native state of iSpinach the fluorophore binding pocket is located between the G-quadruplex and an A-U base pair. The cryo-EM model shows that the bending occurs right at the binding pocket and distorts the G-quadruplex and A-U base pair to occupy the space of the fluorophore. The G-quadruplex is generally preserved, and the main bending seems to occur right at the binding site of the fluorophore. The hinge-like deformation provides an explanation for the mechanical deactivation of the iSpinach aptamer inflicted by the forces of bKL formation in the origami frame.

## DISCUSSION

In this work we have developed a novel RNA device that traps an aptamer in an inactive conformation by embedding it onto an origami frame. Using bKLs to lock the frame, the device was designed to recognize small RNA sequences that trigger the opening of the trap and activation of the aptamer. We observe a cooperative effect on fluorescence when adding both keys, which can be interpreted as Boolean AND gate behavior using an appropriate threshold. Furthermore, because the device uses TMSD for activation it can also be cycled between on and off states. Thus, we have demonstrated an RNA robotic device that can sense, compute and actuate by mechanical force in a reversible manner.

The rate of activation of the Traptamer of 20-30 min is slower than standard RNA strand displacement kinetics [57]. Factors that may contribute to the slow activation kinetics are: (1) The stability of the bKL stems, which was found to correlate to the speed of opening in presence of different keys. (2) Strand displacement into the stable tertiary motif of the kissing loop. (3) Maturation time of the iSpinach aptamer after release from the trap. When testing reversibility of the Traptamer, we observed an even slower activation rate of 80 min and a fast deactivation rate of 10 min. The fast off-rate matches the expected time for a strand displacement and, therefore, the formation of kissing loops and inactivation of iSpinach is likely to happen very fast. The slower on-rate and the faster off-rate can be explained by the fact that internal toeholds tend to slow down the strand displacement process [57]. The speed of activation may be improved by designing less stable bKL stems and by using larger loops to improve toehold binding.

Structural investigation of the 14-14 trap using cryo-EM revealed a bending of the iSpinach aptamer caused by the tension induced on the middle helix due to the formation of the bKLs. Our data suggests that binding sites or active sites may be especially sensitive to mechanical deformation because of the more complex base pairing patterns. This RNA origami framework could be used to study the force required to inactivate the different RNA motifs by changing the number and strength of the kissing loop interactions, effectively tuning the strength of the distortive framework. Similar to DNA origami devices for measuring forces [60-64], the RNA origami framework could be used to measure forces on the nanoscale.

Because RNA origami design is highly modular, the Traptamer can easily be modified to respond to other environmental signals. For example, by using kissing-loop interactions responsive to small molecules [65-67], the Traptamer could be designed to switch upon sensing other molecules and environmental cues. The modularity of the RNA origami architecture also enables the regulation of different functional RNA motifs using similar strategies. We demonstrated this to a limited extent by testing the incorporation of another reporter aptamer, Broccoli [55] into a Traptamer (Fig. S6). By incorporating different fluorescent aptamers and bKL sequences in the origami frames, it may be possible to build multiplex sensor platforms for detecting multiple RNA targets simultaneously. Any other functional RNA motifs with similar configurations (i.e., embedded on a stem) such as protein-binding regions, could be similarly regulated.

Our study of the Traptamer demonstrates an approach to combine sensing, computing and actuation modules to obtain molecular devices that rely on allosteric regulation and large conformational changes for their actuation. By combining different modules, a large variety of RNA robotic devices can be constructed for applications in medicine, synthetic biology, and materials science.

## MATERIALS AND METHODS

### RNA sequence design

RNA origami design methods are explained in Geary et al. 2021 [23]. Briefly, using a standard text editor, the different structural motifs are incorporated into a 2D blueprint. The sequence of the conserved RNA motifs (bKLs, iSpinach aptamer) was constrained, as well as the 3’-end with the primer binding site sequence and the 5’-end with GGA, an optimal initiation sequence for the T7 RNA polymerase. The remaining unconstrained sequence was then designed using the perl script “batch-revolvr.pl” available at https://github.com/esa-lab/ROAD and as web server at https://bion.au.dk/software/rnao-design/. The sequence of the T7 RNA polymerase promoter was introduced upstream of the RNA origami and primers were designed for PCR amplification. Blueprints and sequences can be found in Supplementary note 1.

### Synthesis of DNA templates

The DNA templates for the different RNA designs were produced by PCR amplification of double stranded gene fragments (gBlocks) synthesized by Integrated DNA Technologies (IDT). Amplifications were performed in 100 µl reactions containing 1X Phusion HF buffer (NEB), 1 µM of each primer, 200 µM dNTPs (Invitrogen), 4 ng gBlock template and 1 Unit Phusion High-Fidelity DNA polymerase (NEB). The reaction was subjected to a 2-minute initial denaturation at 98 °C, followed by 30 cycles of: 98 °C for 10 s, 68 °C for 15 s and 72 °C for 10 s, followed by a final extension step at 72 °C of 2 minutes and cooling down to 10 °C. The reaction products were purified using the Macherey-Nagel PCR clean-up kit following the manufacturer’s instructions.

### Production and purification of RNA

RNA was produced by *in vitro* transcription. 500 µl reactions were performed by mixing 3-5 µg of the purified DNA templates, 20 mM MgCl_2_, 10 mM NTPs (2.5 mM each), 10 mM DTT, 40 mM HEPES, 50 mM KCl, 2 mM spermidine and in-house produced T7 RNA polymerase. The reaction was incubated at 37 °C for 3 hours and stopped by adding 2 Units of DNase I (NEB), to digest the DNA templates and incubating at 37 °C for 15 minutes. The reactions were centrifuged at 17,000 RPM for 10 min to pellet the precipitated pyrophosphate. The supernatant was loaded onto a Superose-6 Increase 10/300GL size exclusion column (GE Healthcare/Cytiva) equilibrated with 40 mM HEPES pH 7.5, 50 mM KCl and 5 mM MgCl_2_. The concentration of the co-transcriptionally folded RNA was determined by absorbance measurements at 260 nm on a DeNovix DS-11.

### Fluorescence measurements

Measurements were performed at room temperature (RT, 24°C) on a 384 well plate (Costar), sample volumes of 50 µl contained 100 nM RNA, 500 nM DFHBI-1T and buffer (40 mM HEPES, 50 mM KCl and 5 mM MgCl_2_). 1 µl of Keys were added to specified concentrations (0 to 1µM). (5Z)-5-[(3,5-Difluoro-4-hydroxyphenyl)methylene]-3,5-dihydro-2-methyl-3-(2,2,2-trifluoroethyl)-4H-imidazol-4-one (DFHBI-1T) was purchased from Lucerna Technologies. Fluorescence was monitored using a CLARIOstar Plus multimode microplate reader (BMG LABTECH). Excitation of DFHBI-1T was performed at 470 nm and emission was recorded at 505 nm. For the reversibility experiments, 50 µl containing 100 nM RNA and buffer were mixed. 1 µl of Keys with toehold were added to 500 nM, then 1 µl of Anti-key invaders to 1 µM and finally 1 µl of keys with toehold to 2 µM.

### Cryo-EM sample preparation and data collection

Samples were up-concentrated to ∼ 2mg/ml using Amicon ultra centrifugal filters with a 30 kDa cut-off, spinning the samples at 14000g at RT. ProtoChips Au-FLAT 1.2/1.3 300 mesh grids were glow-discharged for 45 seconds at 15 mA in a Pelco easiGlow prior to sample application. Grids were plunge frozen using a Leica GP2, the sample application chamber was kept at 100% humidity and 21 °C. 3 µl of sample was applied to the grid which was then blotted with a manually calibrated stopping distance onto a double layer of Watman #1 filter paper using a 4 second delay after application, 6 seconds of blot time and 0 seconds of delay after blotting before plunging into liquid ethane at –184 °C.

Data was acquired at 300 keV on a Titan Krios G3i (Thermo Fisher Scientific) equipped with a K3 camera (Gatan/Ametek) and energy filter operated in EFTEM mode using a slit width of 20 eV. Data was collected over a defocus range of -0.5 to -2 micrometers with a targeted dose of 60 e-/Å2. Automated data collection was performed with EPU and the data saved as gain normalized compressed .tiff files (K3) with a pixel size of 0.645 Å/px.

### Cryo-EM single particle analysis data processing

CS-Live was used to apply motion correction, CTF fitting, initial particle picking and an initial *ab initio* model [69]. The 3D volume from CS-live was used for a homogeneous refinement in CryoSPARC V3.2. 50 2D templates were created from this refined volume and templated particle picking was performed using these 50 templates. Particles were then extracted at a pixel size of ∼2.7 Å/px and 3 *ab initio* models were generated using a subset of 30,000 randomly selected particles. A heterogeneous refinement using the three *ab initio* models and all the extracted particles was then performed. At this point we had 1 junk class and 2 classes resembling our RNA origami trap. Another round of heterogeneous refinement was performed using the particles from the 2 good classes and three copies of the best 3D volume as initial search volumes. ∼550,000 particles from the two best classes were then used to start a 5-class *ab initio* model search, followed by heterogeneous refinement into these 5 classes. This resulted in a subsection of 220,000 particles that were reaching the Nyquist limit set by our pixel size, 5.45 Å.

### Model building

Model building was performed in ChimeraX [70, 71]. The different motifs composing the 14-14 trap were manually incorporated to fit into the cryo-EM volume. The crystal structure of iSpinach (PDB: 5OB3) was truncated and fit. The crossovers, helical components and tetraloops were generated using RNAbuild [23], leaving free 5’ and 3’ ends at the crossover junctions, and then manually positioned into the cryo-EM volume. The bKLs were generated by a combination of the NMR structure of the HIV-1 kissing loops (PDB: 2D1B) and a stem loop generated with RNAbuild. Once helical placement was approximately correct, the individual components were joined using the “make bond” command from the ISOLDE [72] add-on to ChimeraX. The resulting PDB was re-numbered using the PDB-Tools pdb-reres program [73] and then the correctly numbered PDB was sequence-corrected in ChimeraX using the swapNA command. This model was then passed through real space refinement (RSR) in Phenix [74-76] (using default parameters, our best refined volume and the resolution supplied by the FSC curve at a 0.143 cutoff) to remove any severe clashes that ISOLDE could not handle. The models were then inspected in ChimeraX and subjected to further refinement using Molecular Dynamics Flexible Fitting (MDFF) with VMD using ISOLDE. A final round of RSR in Phenix was performed to optimize the backbone angles. Validation of the goodness of fit between model and map were performed using the Phenix validation tool [77-79].

## DATA AVAILABILITY

The atomic coordinates for the RNA origami Trap 14-14 RNA have been deposited in the PDB (https://www.rcsb.org/) under the PDB ID XXXX. The volume from the final refinement of our cryo-EM SPA datasets have been deposited to the ePDB under accession codes EMDB-XXXX. Other data are available from the corresponding author upon request.

## AKNOWLEDGEMENTS

We thank Rita Rosendahl and Claus Bus for technical assistance.

## CONTRIBUTION

N. S. V., C. G. and E. S. A. conceptualized the project and designed the experiments. N. S. V. designed the RNAs and performed the fluorescence experiments. E. K. S. M. performed the cryo-EM characterization and analysis. N. S. V., C. G. and E. S. A. wrote the manuscript.

## ETHICS DECLARATION

The authors declare no competing interests.

