## Supplementary Information for "An RNA origami robot that traps and releases a fluorescent aptamer"

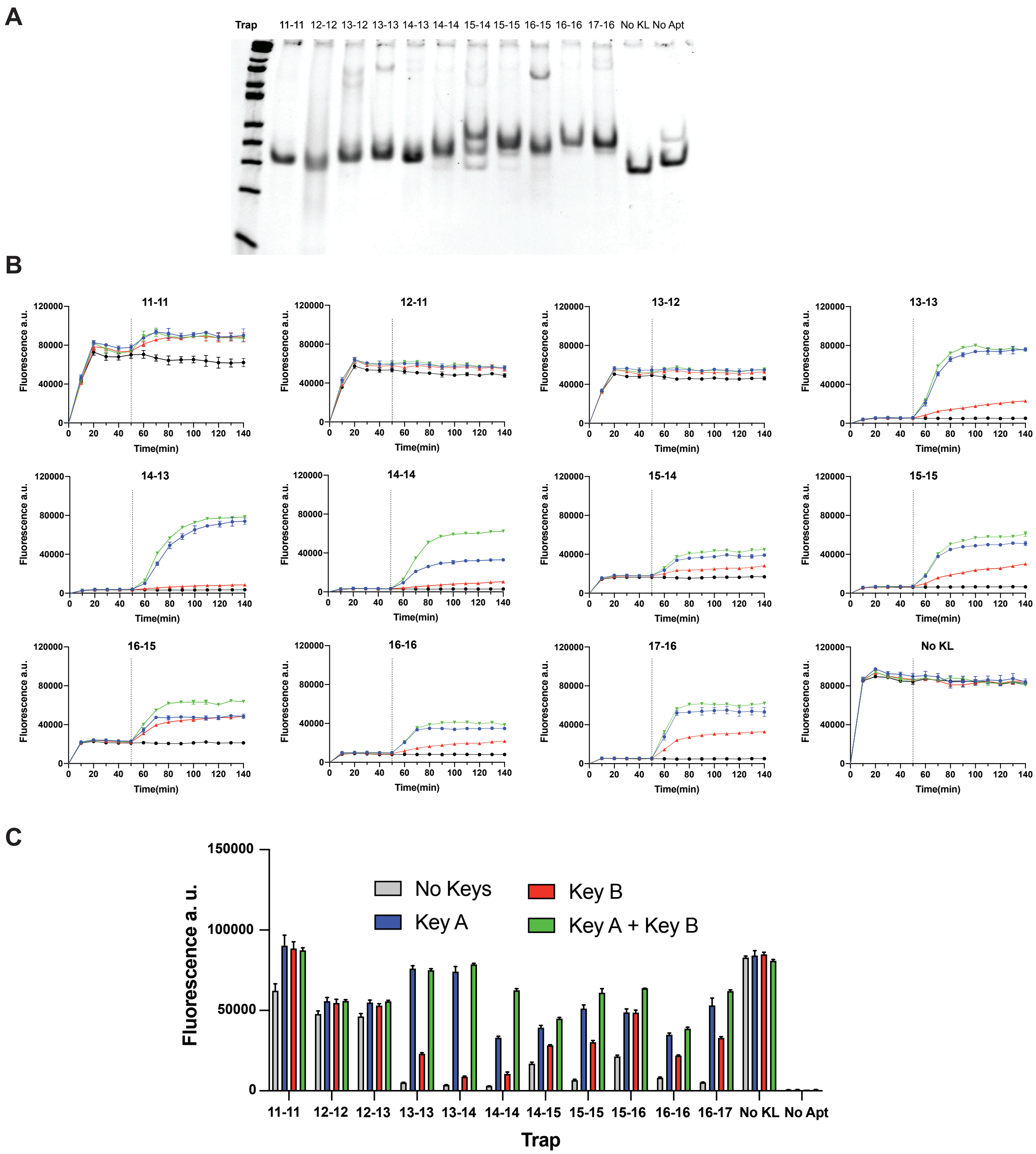

**Figure S1. Design screening for a co-transcriptionally folded RNA mechanical trap.**
**A.** 8% Native PAGE gel showing the state of the different designs after SEC purification. **B.** Design screening. Monitored fluorescence over time of the different designs. The different RNAs were produced and purified to a final concentration of 100 nM. The fluorophore DFHBI-1T (1 µM) was added after the first measurement and 5X (500 nM) single stranded RNA keys were added after 50 minutes (black dashed line). Data corresponds to 3 technical replicates, error bars represent mean ± SD. **C.** Fluorescence observed at 140 mins (85 minutes upon addition og the RNA keys) for the different designs.

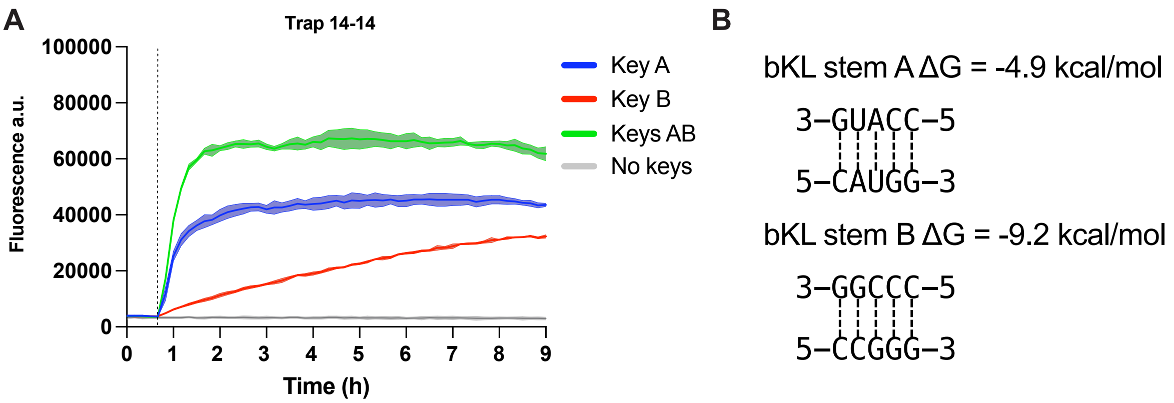

**Figure S2. Opening of Trap-14-14 and stabilities over extended time period.**
**A.** Monitored fluorescence over 9 hours of trap 14-14 (100 nM) in solution with DFHBI-1T (500 nM) after addition of RNA keys (500 nm) at RT (dashed line indicates key addition). Data represents 3 technical replicates ± SD. **B.** Free energies of the bKL stems calculated with Nupack.

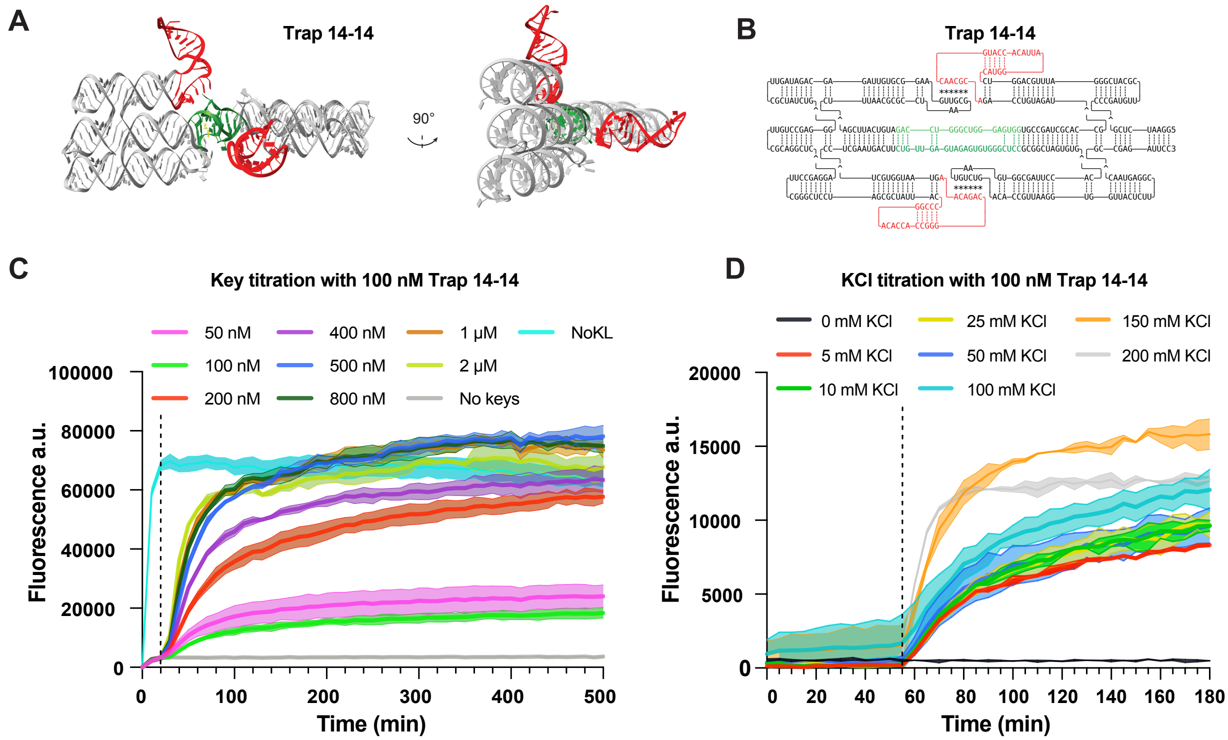

**Figure S3. Model and properties of the Trap-14-14 design.**

**A.** Computer-generated 3D model with RNAbuild [1]. The iSpinach aptamer is depicted in green, bKLs in red and DFHBI-1T ligand in yellow. **B.** RNA sequence text blueprint. The iSpinach motif is highlighted in green and the bKLs in red. **C.** ssRNA key titration. Monitored fluorescence over time of Trap-14-14 RNA (100 nM) in complex with DFHBI-1T (1 µM), different concentrations of ssRNAs were added after the first fluorescence measurement. The No kissing loops (NoKL) positive control is shown in cyan blue. Data corresponds to 3 technical replicates, error bars represent mean ± SD. **D.** KCl titration. Monitored fluorescence over time of Trap-14-14 (100 nM) in buffers with different KCl concentrations. Data corresponds to 3 technical replicates, error bars represent mean ± SD.

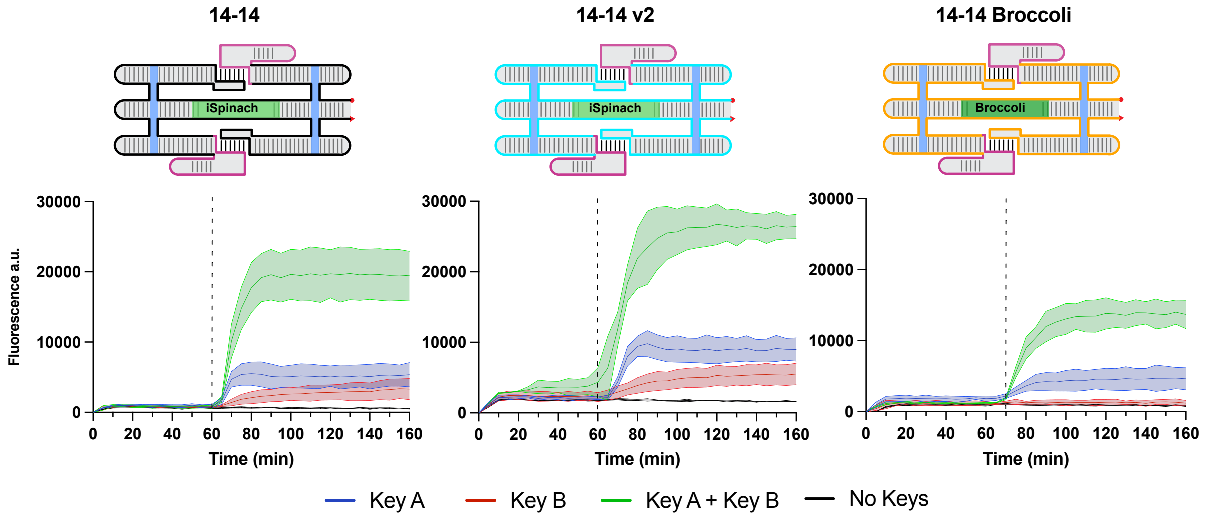

**Figure S4. Function of Trap 14-14 with different sequence designs and Broccoli aptamer.**

Observed fluorescence for the 14-14 design variants over time. 14-14 v2 has a different scaffold sequence (blue line). 14-14 Broccoli has the Broccoli aptamer and a different scaffold sequence (orange line). DFHBI-1T (500 nM) keys (500 nM) were added to 100 nM RNA after 60 minutes (dashed line). Data corresponds to 3 technical replicates; error bars represent mean ± SD.

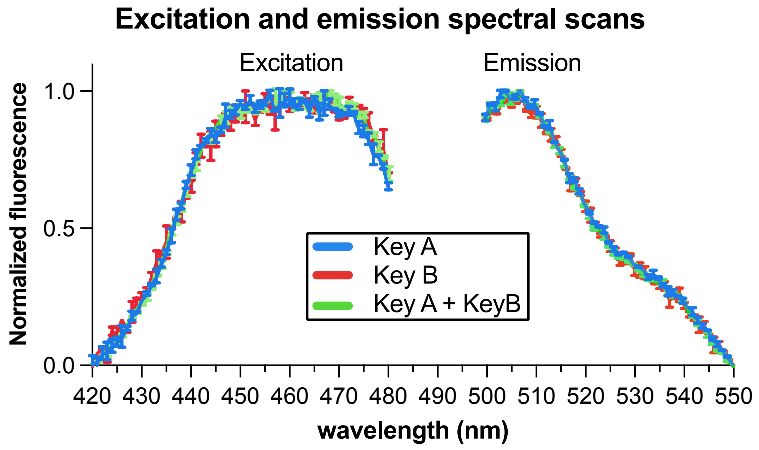

**Figure S5. No effect of key addition on excitation and emission spectra.**
Excitation and emission spectra of DFHBI-1T (500 nM) in complex with the Trap 14-14 (100 nM) at different key conditions (500 nM).

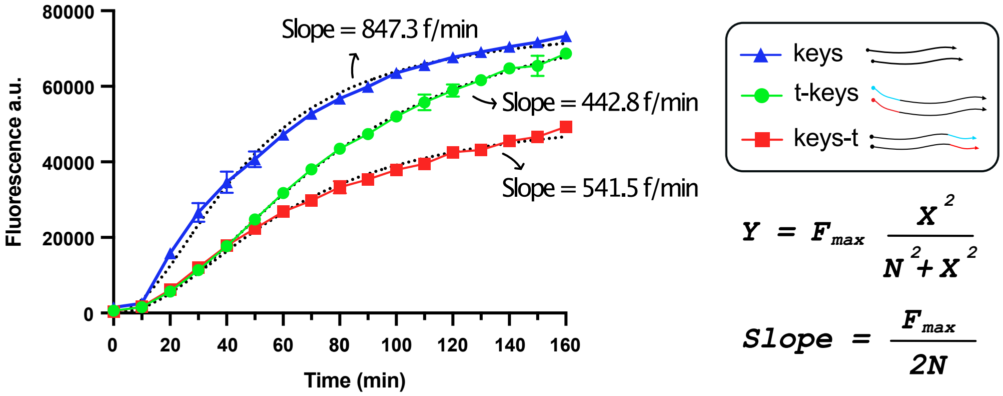

**Figure S6. Effect of adding toeholds the ends of RNA keys.**

Fluorescence activation of 100 nM Trap 14-14 after addition of keys (500 nM) with no toehold or toehold placed at 5'- (t-keys) or 3'- (keys-t) ends (500 nM). Data represents 3 technical replicates ± SD. $F_{max}$ represents the maximum fluorescence and *N* represents the inflection point, i.e., time point (x) at which fluorescence has its maximum increase. Model fit $R^2$ = 0.993 for keys, $R^2$ = 0.998 for t-keys and $R^2$ = 0.993 for keys-t.

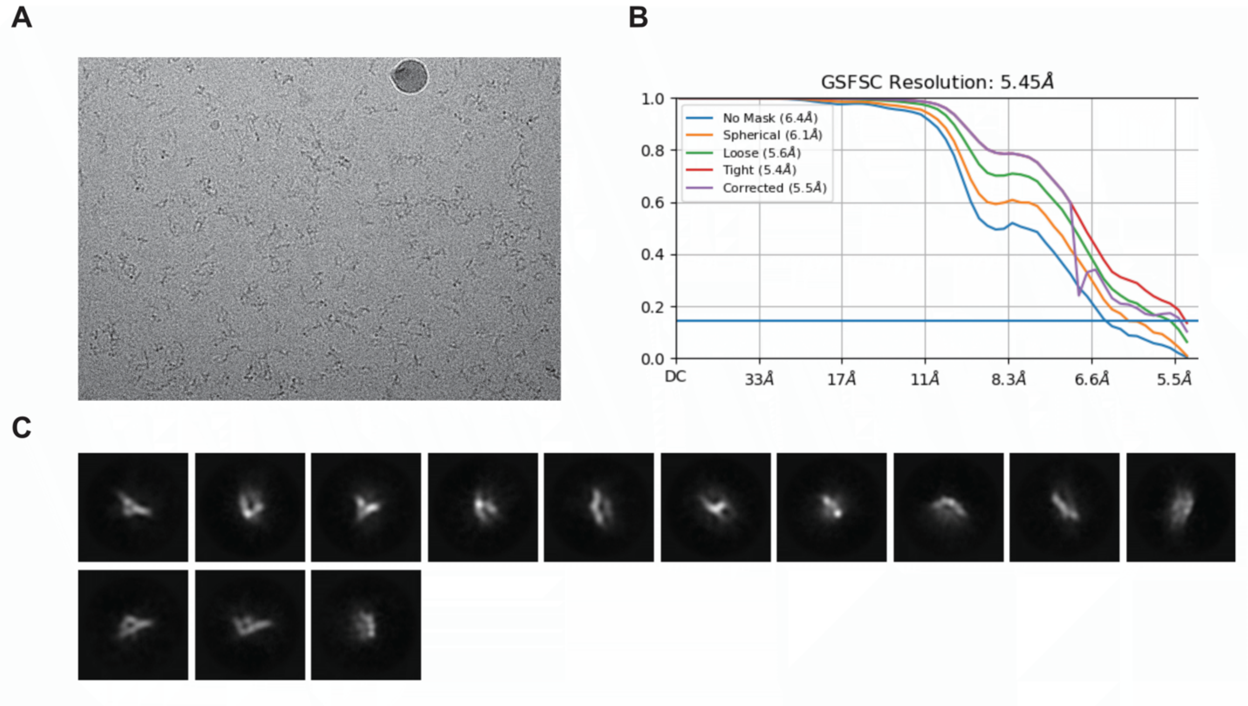

**Figure S7. Cryo-EM analysis of Trap-14-14 with no keys.**
**A.** Example cryo-EM micrograph from the locked Trap-14-14 dataset. **B.** Gold-standard Fourier Shell Correlation for the locked state of Trap-14-14. **C.** 2D Class. Averages from the final particle stack of the locked Trap-14-14 dataset.

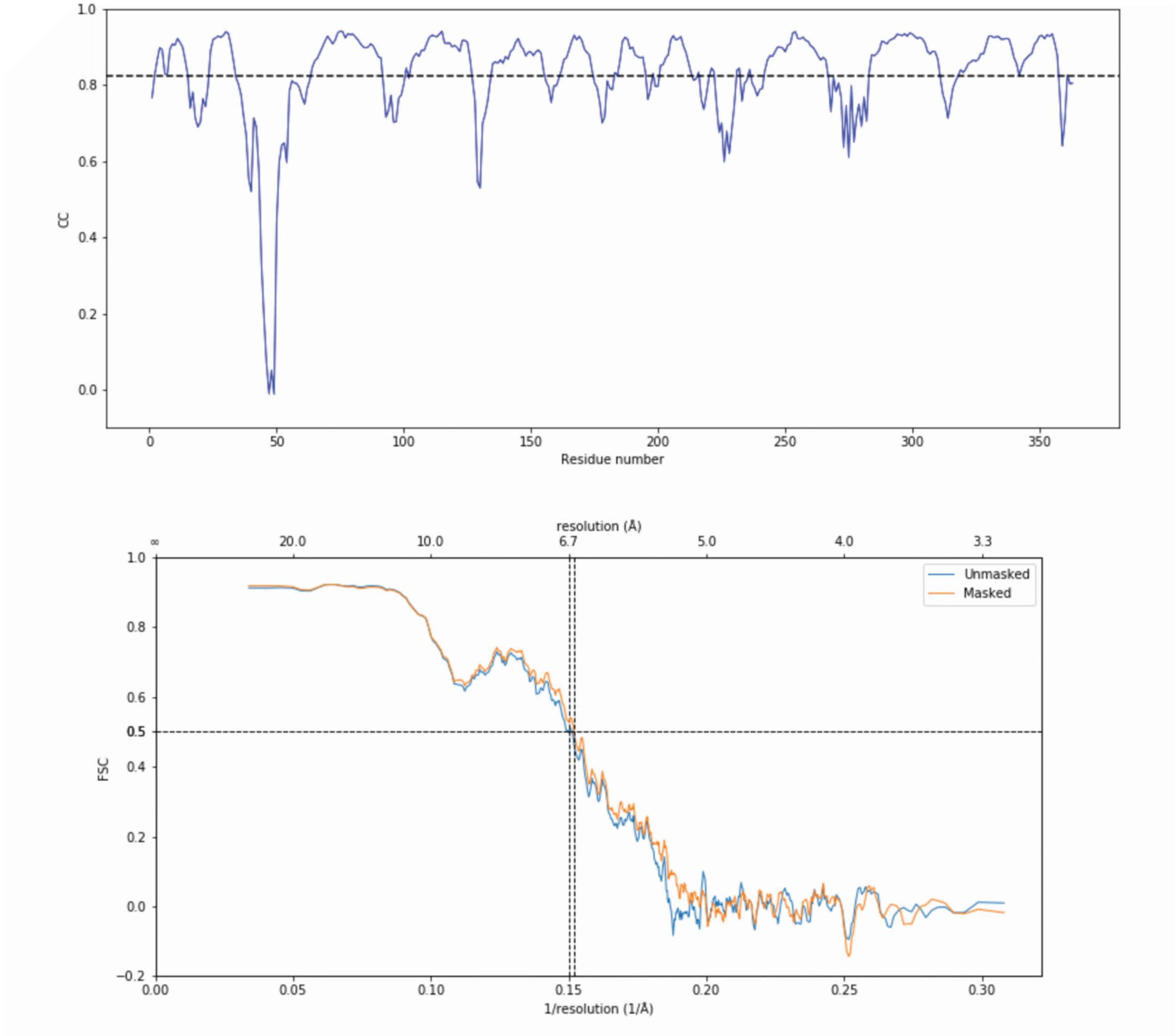

**Figure S8. Cross Correlation and Fourier Shell Correlation.**
Per residue Cross Correlation (top) and Fourier Shell Correlation (bottom) between atomic model and cryo-EM map for the trap 14-14 with no keys after phenix RSR.

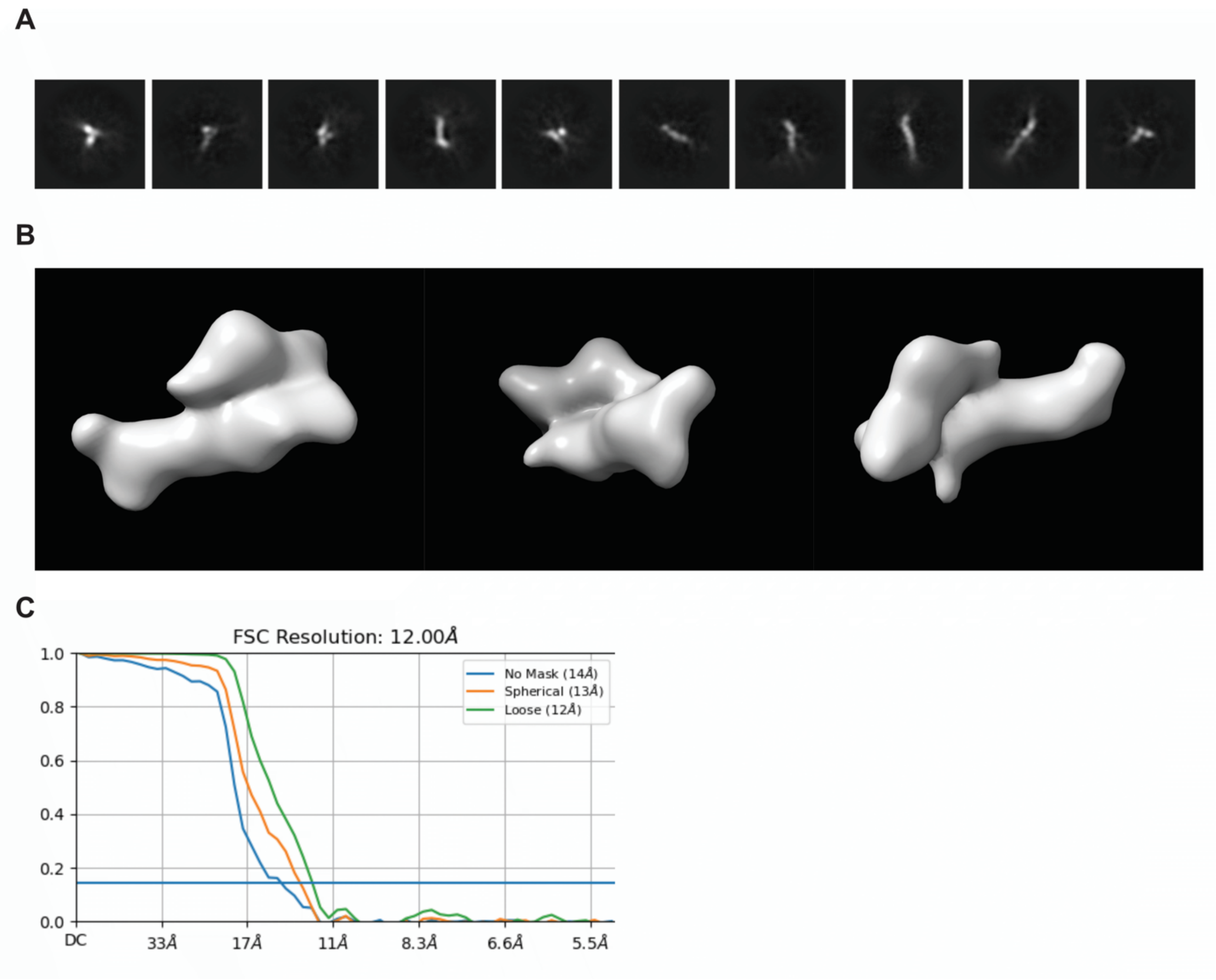

**Figure S9. Cryo-EM analysis of Trap-14-14 upon addition of keys.**
**A.** 2D Class averages from the. Final particle stack of the open Trap-14-14. dataset. **B.** Three views of the Trap-14-14 device with keys reconstruction. **C.** Gold-Standard Fourier Shell Correlation for the open state of Trap-14-14.

### Table S1. RNA blueprints and sequences

| No Kissing loops control (NoKL) |
|---|
| 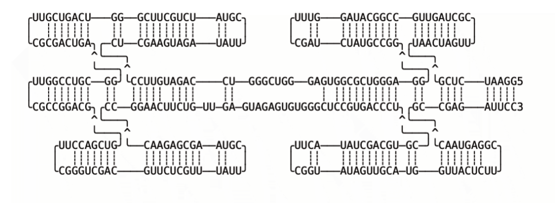<br>= |
| GGAAUCUCGUAACUAGUUCGCUAGUUGCCGGCAUAGGUUUCGAUCUAUGCCGGGCAGGGUCGCGGUGAGGGUCGGGUCCAGAUGUUCCCUCGAAGUAGAUA<br>UUCGUAUCUGCUUCGGGUCAGUCGUUCGCGACUGAGGCGUCCGGUUCGCCGGACGGUCGACCUUCGGGUCGACGUUCUCGUUUAUUCGUAAGCGAGAACCC<br>GGAACUUCUGUUGAGUAGAGUGUGGGCUCCGUGACCCUCGUGCAGCUAUACUUCGGUAUAGUUGCAUGGUUACUCUUCGGAGUAACGCCGAGAUUCC |

| No Aptamer control (NoApt) |
|---|
| 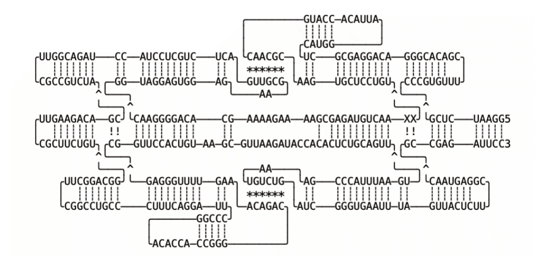 |
| GGAAUCUCGCCCGUGUUUCGACACGGGACAGGAGCGCUCAUGGAUUACACCAUGCAACGCAAGUGCUCCUGUGCAACUGUAGAGCGAAAAGAAAAGCACAG<br>GGGAACGGUAGGAGUGGAGAAGCGUUGACUCUGCUCCUACCUAGACGGUUCGCCGUCUACGACAGAAGUUCGCUUCUGUGGCAGGCUUCGGCCUGCCCUUU<br>CAGGAUUCCCGGACACCACCGGGCAGACAAAGUUUUGGGAGCGGUUCCACUGUAAGCGUUAAGAUACCACACUCUGCAGUUUGAAUUUACCCGAAAUGUCU<br>GAUCGGGUGAAUUUAGUUACUCUUCGGAGUAACGCCGAGAUUCC |

| Trap-11-11 |
|---|
| 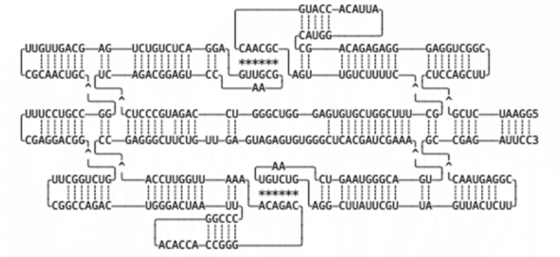 |
| GGAAUCUCGCUCCAGCUUCGGCUGGAGGGAGAGACAGCCAUGGAUUACACCAUGCAACGCAGUUGUCUUUUCGCUUUCGGUCGUGUGAGGGUCGGGUCCAG<br>AUGCCCUCUCAGACGGAGUCCAAGCGUUGAGGACUCUGUCUGAGCAGUUGUUCGCAACUGCGGCCGUCCUUUCGAGGACGGGUCUGGCUUCGGCCAGACUG<br>GGACUAAUUCCCGGACACCACCGGGCAGACAAAAUUGGUUCCACCGAGGGCUUCUGUUGAGUAGAGUGUGGGCUCACGAUCGAAAUGACGGGUAAGUCAAU<br>GUCUGAGGCUUAUUCGUUAGUUACUCUUCGGAGUAACGCCGAGAUUCC |

### Trap-12-12

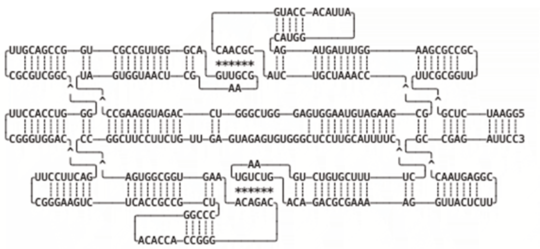

GGAAUCUCGUUCGCGGUUCGCCGCGAAGGUUUAGUAGACAUGGAUUACACCAUGCAACGCAUCUGCUAAACCGCGAAGAUGUAAGGUGAGGGUCGGGUCCA
GAUGGAAGCCUAGUGGUAACUCGAAGCGUUGACGGGUUGCCGCUGGCCGACGUUCGCGUCGGCGGGUCCACCUUCGGGUGGACGACUUCCUUCGGGAAGUC
UCACCGCCGCUCCCGGACACCACCGGGCAGACAAAGUGGCGGUGACCGGCUUCCUUCUGUUGAGUAGAGUGUGGGCUCCUUGCAUUUUCCUUUUCGUGUCU
GAAUGUCUGACAGACGCGAAAAGGUUACUCUUCGGAGUAACGCCGAGAUUCC

### Trap-13-12

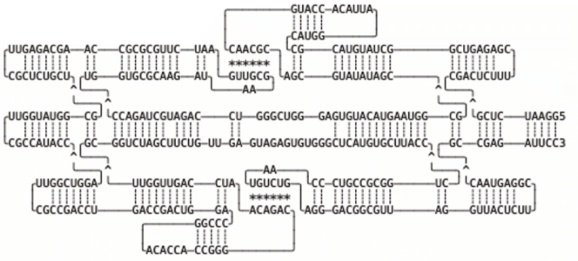

GGAAUCUCGCGACUCUUUCGAGAGUCGGCUAUGUACGCCAUGGAUUACACCAUGCAACGCAGCGUAUAUAGCGCGGUAAGUACAUGUGAGGGUCGGGUCCA
GAUGCUAGACCUGGUGCGCAAGAUAAGCGUUGAAUCUUGCGCGCCAAGCAGAGUUCGCUCUGCUGCGGUAUGGUUCGCCAUACCAGGUCGGUUCGCCGACC
UGACCGACUGGACCCGGACACCACCGGGCAGACAAUCCAGUUGGUUGCGGUCUAGCUUCUGUUGAGUAGAGUGUGGGCUCAUGUGCUUACCCUGGCGCCGU
CCCAAUGUCUGAGGGACGGCGUUAGGUUACUCUUCGGAGUAACGCCGAGAUUCC

### Trap-13-13

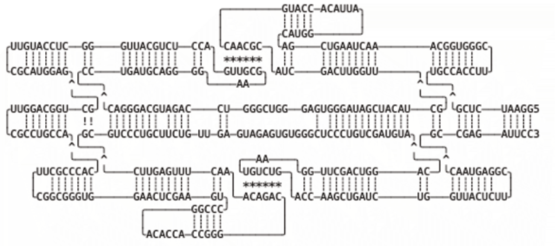

GGAAUCUCGUGCCACCUUCGGGUGGCAAACUAAGUCGACAUGGAUUACACCAUGCAACGCAUCGACUUGGUUGCUACAUCGAUAGGGUGAGGGUCGGGUCC
AGAUGCAGGGACCCUGAUGCAGGGGAAGCGUUGACCUCUGCAUUGGGCUCCAUGUUCGCAUGGAGGCUGGCAGGUUCGCCUGCCACACCCGCUUCGGCGGG
UGGAACUCGAAGUCCCGGACACCACCGGGCAGACAAACUUUGAGUUCGCGUCCCUGCUUCUGUUGAGUAGAGUGUGGGCUCCCUGUCGAUGUACAGGUCAG
CUUGGAAUGUCUGACCAAGCUGAUCUGGUUACUCUUCGGAGUAACGCCGAGAUUCC

### Trap-14-13

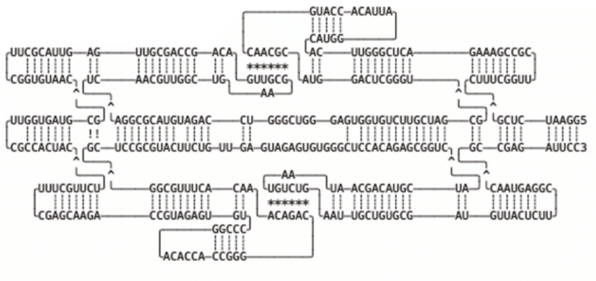

GGAAUCUCGCUUUCGGUUCGCCGAAAGACUCGGGUUCACAUGGAUUACACCAUGCAACGCAUGGACUCGGGUGCGAUCGUUCUGUGGUGAGGGUCGGGUCC
AGAUGUACGCGGAUCAACGUUGGCUGAAGCGUUGACAGCCAGCGUUGAGUUACGCUUCGGUGUAACGCGUAGUGGUUCGCCACUACUCUUGCUUUCGAGCA
AGACCGUAGAGUGUCCCGGACACCACCGGGCAGACAAACACUUUGCGGGCUCCGCGUACUUCUGUUGAGUAGAGUGUGGGCUCCACAGAGCGGUCAUCGUA
CAGCAAUAAUGUCUGAAUUGCUGUGCGAUGUUACUCUUCGGAGUAACGCCGAGAUUCC

### Trap-14-14

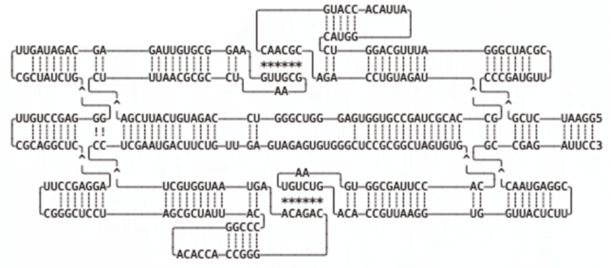

GGAAUCUCGCCCGAUGUUCGCAUCGGGAUUUGCAGGUCCAUGGAUUACACCAUGCAACGCAGACCUGUAGAUGCCACGCUAGCCGUGGUGAGGGUCGGGUC
CAGAUGUCAUUCGACUUUAACGCGCCUAAGCGUUGAAGGCGUGUUAGAGCAGAUAGUUCGCUAUCUGGGGAGCCUGUUCGCAGGCUCAGGAGCCUUCGGGC
UCCUAGCGCUAUUACCCCGGACACCACCGGGCAGACAAGUAAUGGUGCUCCUCGAAUGACUUCUGUUGAGUAGAGUGUGGGCUCCGCGGCUAGUGUGCACC
UUAGCGGUGAAUGUCUGACACCGUUAAGGUGGUUACUCUUCGGAGUAACGCCGAGAUUCC

### Trap-15-14

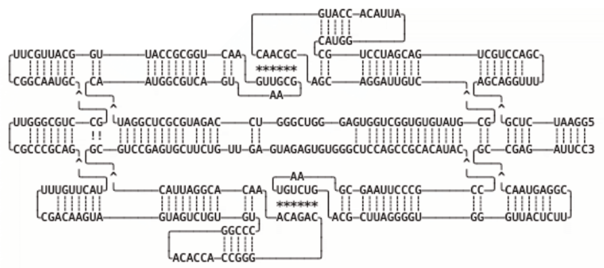

GGAAUCUCGAGCAGGUUUCGACCUGCUGACGAUCCUGCCAUGGAUUACACCAUGCAACGCAGCAGGAUUGUCGCGUAUGUGUGGCUGGUGAGGGUCGGGUC
CAGAUGCGCUCGGAUCAAUGGCGUCAGUAAGCGUUGAACUGGCGCCAUUGGCAUUGCUUCGGCAAUGCGCCUGCGGGUUCGCCCGCAGUACUUGUUUCGAC
AAGUAGUAGUCUGUGUCCCGGACACCACCGGGCAGACAAACACGGAUUACGCGUCCGAGUGCUUCUGUUGAGUAGAGUGUGGGCUCCAGCCGCACAUACCC
GCCCUUAAGCGAAUGUCUGACGCUUAGGGGUGGGUUACUCUUCGGAGUAACGCCGAGAUUCC

### Trap-15-15

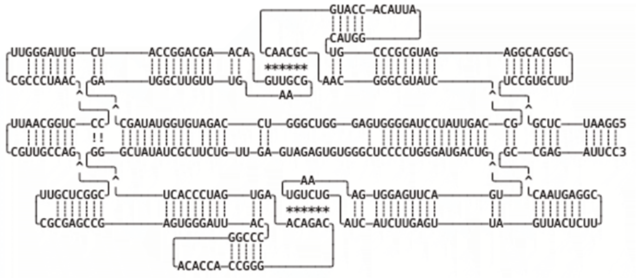

GGAAUCUCGUCCGUGCUUCGGCACGGAGAUGCGCCCGUCAUGGAUUACACCAUGCAACGCAACGGGCGUAUCGCCAGUUAUCCUAGGGGUGAGGGUCGGGU
CCAGAUGUGGUAUAGCGAUGGCUUGUUUGAAGCGUUGACAAGCAGGCCAUCGUUAGGGUUCGCCCUAACCCCUGGCAAUUCGUUGCCAGCGGCUCGUUCGC
GAGCCGAGUGGGAUUACCCCGGACACCACCGGGCAGACAAGUGAUCCCACUGGGCUAUAUCGCUUCUGUUGAGUAGAGUGUGGGCUCCCCUGGGAUGACUG
UGACUUGAGGUGAAAUGUCUGAUCAUCUUGAGUUAGUUACUCUUCGGAGUAACGCCGAGAUUCC

### Trap-16-15

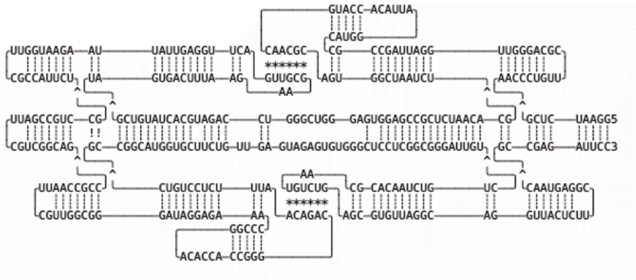

GGAAUCUCGAACCCUGUUCGCAGGGUUGGAUUAGCCGCCAUGGAUUACACCAUGCAACGCAGUGGCUAAUCUGCACAAUCUCGCCGAGGUGAGGGUCGGGU
CCAGAUGCACUAUGUCGUAGUGACUUUAAGAAGCGUUGACUUGGAGUUAUUAAGAAUGGUUCGCCAUUCUGCCUGCCGAUUCGUCGGCAGCCGCCAAUUCG
UUGGCGGGAUAGGAGAAACCCGGACACCACCGGGCAGACAAUUUCUCCUGUCGCCGGCAUGGUGCUUCUGUUGAGUAGAGUGUGGGCUCCUCGGCGGGAUU
GUCUGUCUAACACGCAAUGUCUGAGCGUGUUAGGCAGGUUACUCUUCGGAGUAACGCCGAGAUUCC

### Trap-16-16

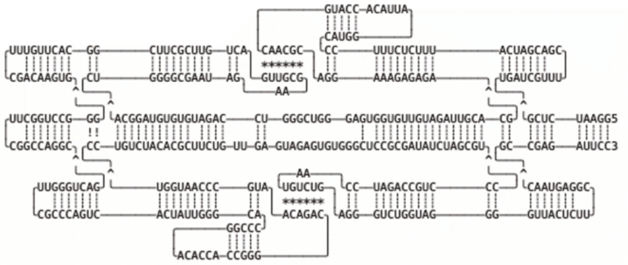

GGAAUCUCGUGAUCGUUUCGACGAUCAUUUCUCUUUCCCAUGGAUUACACCAUGCAACGCAGGAAAGAGAGAGCACGUUAGAUGUUGUGGUGAGGGUCGGG
UCCAGAUGUGUGUAGGCACUGGGGCGAAUAGAAGCGUUGACUGUUCGCUUCGGCACUUGUUUCGACAAGUGGGGCCUGGCUUCGGCCAGGCGACUGGGUUC
GCCCAGUCACUAUUGGGCACCCGGACACCACCGGGCAGACAAUGCCCAAUGGUCCUGUCUACACGCUUCUGUUGAGUAGAGUGUGGGCUCCGCGAUAUCUA
GCGUCCCUGCCAGAUCCAAUGUCUGAGGGUCUGGUAGGGGUUACUCUUCGGAGUAACGCCGAGAUUCC

| Trap-17-16 |
| --- |
| 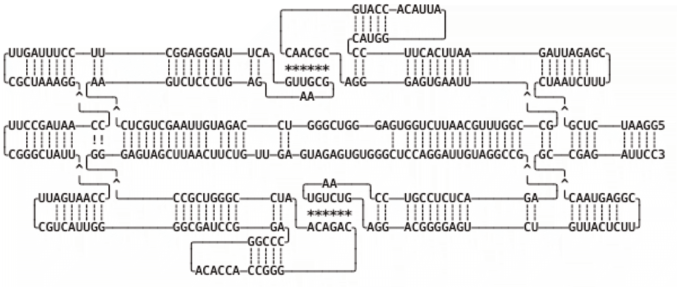 |
| GGAAUCUCGCUAAUCUUUCGAGAUUAGAAUUCACUUCCCAUGGAUUACACCAUGCAACGCAGGGAGUGAAUUGCCGGUUUGCAAUUCUGGUGAGGGUCGGG UCCAGAUGUUAAGCUGCUCAAGUCUCCCUGAGAAGCGUUGACUUAGGGAGGCUUCCUUUAGUUCGCUAAAGGCCAAUAGCCUUCGGGCUAUUCCAAUGAUU CGUCAUUGGGGCGAUCCGGACCCGGACACCACCGGGCAGACAAUCCGGGUCGCCGGGAGUAGCUUAACUUCUGUUGAGUAGAGUGUGGGCUCCAGGAUUGU AGGCCGAGACUCUCCGUCCAAUGUCUGAGGACGGGGAGUCUGUUACUCUUCGGAGUAACGCCGAGAUUCC |

| Trap-14-14 v2 |
| --- |
| 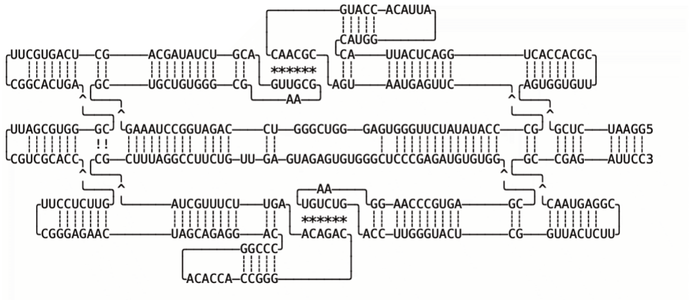 |
| GGAAUCUCGAGUGGUGUUCGCACCACUGGACUCAUUACCAUGGAUUACACCAUGCAACGCAGUAAUGAGUUCGCCCAUAUAUCUUGGGUGAGGGUCGGGUC CAGAUGGCCUAAAGGCUGCUGUGGGCGAAGCGUUGACGUCUAUAGCAGCUCAGUGCUUCGGCACUGACGGGUGCGAUUCGUCGCACCGUUCUCCUUCGGGA GAACUAGCAGAGGACCCCGGACACCACCGGGCAGACAAGUUCUUUGCUACGCUUUAGGCCUUCUGUUGAGUAGAGUGUGGGCUCCCGAGAUGUGUGGCGAG UGCCCAAGGAAUGUCUGACCUUGGGUACUCGGUUACUCUUCGGAGUAACGCCGAGAUUCC |

| Trap-14-14 Broccoli |
| --- |
| 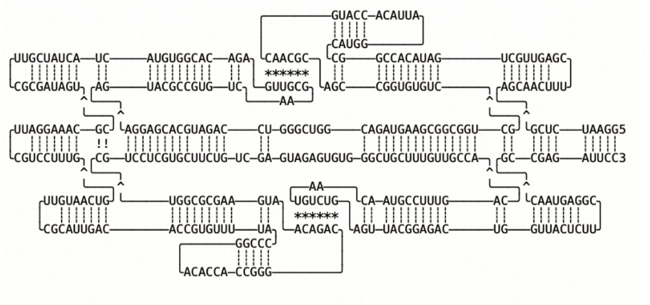 |
| GGAAUCUCGAGCAACUUUCGAGUUGCUGAUACACCGGCCAUGGAUUACACCAUGCAACGCAGCCGGUGUGUCGCUGGCGGCGAAGUAGACGGUCGGGUCCA GAUGCACGAGGAAGUACGCCGUGUCAAGCGUUGAGACACGGUGUACUACUAUCGUUCGCGAUAGUCGCAAAGGAUUCGUCCUUUGGUCAAUGUUCGCAUUG ACACCGUGUUUUACCCGGACACCACCGGGCAGACAAUGAAGCGCGGUCGUCCUCGUGCUUCUGUCGAGUAGAGUGUGGGCUGCUUUGUUGCCACAGUUUCC GUAACAAUGUCUGAGUUACGGAGACUGGUUACUCUUCGGAGUAACGCCGAGAUUCC |

**Table S2. PCR primers and RNA keys.**

**PCR Primers (5' to 3')**

| Forward |
| --- |
| CAGACTTCTTACGTGATCCATCCG |
| Reverse |
| GGAATCTCGGCGTTACTCCG |

**RNA keys (5' to 3')**

| Key A |
| --- |
| GCGUUGCAUGGUGUAAU |
| Key B |
| UGUCUGCCCGGUGGUGU |
| Key C |
| AACCUAACUCAAUCUCU |
| Key D |
| UAACAAAACAGACAAAG |
| Key A with 3' toehold |
| GCGUUGCAUGGUGUAAU-ACGUCG |
| Key B with 3' toehold |
| UGUCUGCCCGGUGGUGU-UGAAGC |
| Anti-key t-A* |
| CGACGUAUUACACCAUGCAACGC |
| Anti-key t-B* |
| GCUUCAACACCACCGGGCAGACA |
| Key A with 5' toehold |
| GUCCAC-GCGUUGCAUGGUGUAAU |
| Key B with 5' toehold |
| CGAAGU-UGUCUGCCCGGUGGUGU |
| Negative control invader E |
| CUGCACAGAACGGGAUUCUUUCA |
| Negative control invader F |
| GGGAAGGUGUUUGCUGGGAGUGA |
